# Thriving of hyperthermophilic microbial communities from a deep-sea sulfidic hydrothermal chimney under electrolithoautotrophic conditions with nitrate as electron acceptor

**DOI:** 10.1101/2021.03.26.437165

**Authors:** G. Pillot, S. Davidson, L. Shintu, L. Tanet, Y. Combet-Blanc, A. Godfroy, P. Bonin, P.-P. Liebgott

## Abstract

Recent studies have shown the presence of an abiotic electrical current across the walls of deep-sea hydrothermal chimneys, allowing the growth of electroautotrophic microbial communities. To understand the role of the different phylogenetic groups and metabolisms involved, this study focused on an electrotrophic enrichment, with nitrate as electron acceptor. The biofilm density, the community composition, the organic products released, and the electrical consumption were monitored by FISH confocal microscopy, qPCR, Metabarcoding, MNR and potentiostat measurements. A statistic analysis by PCA showed the correlation between the different parameters in 3 distinct temporal phases. The *Archaeoglobales* have been shown to play a key role in the development of the community, as first colonizers and producing pyruvate, therefor used as organic source for heterotrophs. Some *Thermococcales* showed the ability to perform electrofermentation of this pyruvate into acetate and H_2_. Finally, through subcultures of the community, we showed the development of a larger biodiversity over time. This observed phenomenon could explain the biodiversity development in hydrothermal context where energy sources are transient and unstable.

## Introduction

Deep-sea hydrothermal vents, discovered for the first time in 1977, represent complex ecosystem sheltering extremophilic life forms [1, 2]. These hydrothermal chimneys result from the infiltration of seawater in the seabed, heated by an underlying magma chamber and reacting with surrounding minerals to produce a hot hydrothermal fluid (ca. 300-400°C) rich in minerals and reduced compounds (H_2_, H_2_S, CH_4_, CO_2_). This hot reduced fluid under pressure will move back to the surface to precipitate in contact with the cold (ca. 2°C) and oxidized surrounding seawater. These reactions form an area of intense mixing and severe thermal and chemical gradients allowing the development of a complex and extremophilic biosphere. Because of the high temperature of the hydrothermal fluid, considered as sterile [3], the question of first colonizers of these newly formed hydrothermal chimneys arises [3–5]. It is now accepted that, in the absence of organic matter and light, only chemolithoautotrophs coming from seawater or earth’s crust can grow in the first stage of colonization. The colonizers would use the oxidation of reduced compounds from the hydrothermal fluid as an energy source and CO_2_ as a carbon source to produce organic matter. Some studies have attempted to study the microbial diversity in a newly formed chimney through the formation of a new chimney in mineral chambers and sampling units or on thermocouples placed in the hydrothermal fluid [4, 5]. They highlighted the presence of *Desulfurococcaceae*, mainly members of the genus *Ignicoccus*, and its symbiont *Nanaoarchaeum* in addition to *Thermococcus* sp. and *Methanocaldococcus* spp. This primary production of organic matter would subsequently support the growth of large diversity of heterotrophs through the complex trophic chain and microbial interactions [6, 7]. Besides the enrichments of hyperthermophilic *Archaea*, recent study postulated that colonization of a newly erupted black smoker occurs by hyperthermophiles [8]. Two hyperthermophilic species - from *Thermococcales* and *Methanococcales* - have been shown to swim to mineral surface, scanning the surface to find optimal temperature condition, to finally adhere directly on it through mechanisms that could be similar to those of the bacterial pili IV.

Most of the methods used to enrich these first colonizers in the laboratory have postulated the use of reduced compounds from the hydrothermal fluid as substrates. However, recent studies have shown the presence of an abiotic electrical current inside the chimney wall produced by chemical oxidation of hydrogen sulfide from hydrothermal fluid coupled with oxygen reduction from seawater [9]. It has been hypothesized that this electrical current could be used as a direct energy source by electroactive microbes. These microbes, extensively studied for 20 years, have the ability to perform external electron transfer towards or from conductive support for their metabolism [10]. Microorganisms able to use direct electron flow as an energy source are called electrotrophs. They can take up directly electrons from the cathode, the electron donor, in a Microbial Electrochemical Systems (MES) for their energy and transfer it into a terminal electron acceptor. In the literature, many studies have aimed to enrich these electrotrophs within microbial communities sampled from various ecosystems [11–14] or to axenically grow them [15, 16] Recently, we have successfully enriched hyperthermophilic electrotrophs from hydrothermal vent samples [17]. The only presence of CO_2_ as carbon source, an electron acceptor, and an abiotic electron flow fed by the MES cathode as energy source were sufficient to grow “electro-autotrophic” microorganisms from hydrothermal chimneys. The community obtained in the electrotrophic biofilm was mainly composed of archaea from *Archeoglobales* and *Thermococcales* with the specific enrichment of other phylogenetic groups depending on the electron acceptor used. This finding leads to raise some questions such as: is this first primary production enough to drive the development of more complex biodiversity through the food web and microbial interactions? What is the importance of the electro-autotrophic ability within the theory of the origin of life in a deep hydrothermal environment? In this latter context, nitrate was used as the electron acceptor, for its hypothetical central role in the theory of the origin of life in hydrothermal context[18].

The objective of this work was to study the evolution of the composition of the hyperthermophilic microbial community and organic products over time, to identify the first colonizers and the metabolisms involved. This microbial community was cultured in electrolithoautotrophic condition with polarized cathode as electron donor and nitrate as the terminal electron acceptor. Thus, current density related to electrotrohic growth and nitrate consumption were monitored, along with measurements of organic and gaseous compounds produced as well as the quantification by qPCR of dominant phylogenetic orders. Afterwards, we studied if this electrotrophic community could lead to a more complex ecosystem through two subsequent subcultures in MES.

## Materials and Methods

### Sample collection and preparation

A hydrothermal chimney was collected on the Capelinhos site on Lucky Strike hydrothermal field (37°17.0’N, MAR) during the MoMARsat cruise in 2014 (doi: 10.17600/14000300) led by IFREMER (France) onboard R/V *Pourquoi Pas?* [19]. The sample (PL583-8) was collected and prepared in anaerobic conditions as previously described to inoculate a Microbial Electrochemical Systems [20].

### Electrotrophic enrichment in a microbial electrochemical system

A Microbial Electrochemical System (MES) was filled with 1.5 L of sterile mineral medium as previously described [21] with 0.1 g/l of Cysteine-HCl and supplemented with 4mM Nitrate, continuously flushed with N_2_:CO_2_ (90:10, 100 ml/min) to anaerobic condition and set at 80°C and pH 6.0 throughout platform monitoring. The working electrode (WE) composed of 20 cm^2^ of carbon cloth was poised at −590mV vs SHE using SP-240 potentiostats and EC-Lab software (BioLogic, France). The WE served as the sole electron donor for electrotrophs development. The system was inoculated with 8 g of the crushed chimney in anaerobic conditions. Current consumption was monitored via the chronoamperometry method with current density and counter-electrode potential measurements taken every 10 s.

To evaluate the enrichment of biodiversity over time on the electrode (WE) and in the liquid medium (LM), three successive cultures were performed. For the first culture, shown in Figure 1, a fraction of a crushed chimney from the Capelinhos site was used to inoculate the MES. After 25 days of incubation, the electrode (C1-WE) and the liquid medium (C1-LM) were harvested. An open circuit potential (OCP) control was performed in the same conditions. For the second culture, a new MES with sterile electrode was inoculated with 100 ml of the C1-LM in a fresh mineral medium. The electrode (C2-WE) and liquid media (C2-LM) were harvested after enrichment. The third culture was performed with inoculation from 100 ml of C2-LM. Abiotic control without inoculation didn’t show current consumption or nitrate depletion during the same time of the experiment.

**Figure 1:**
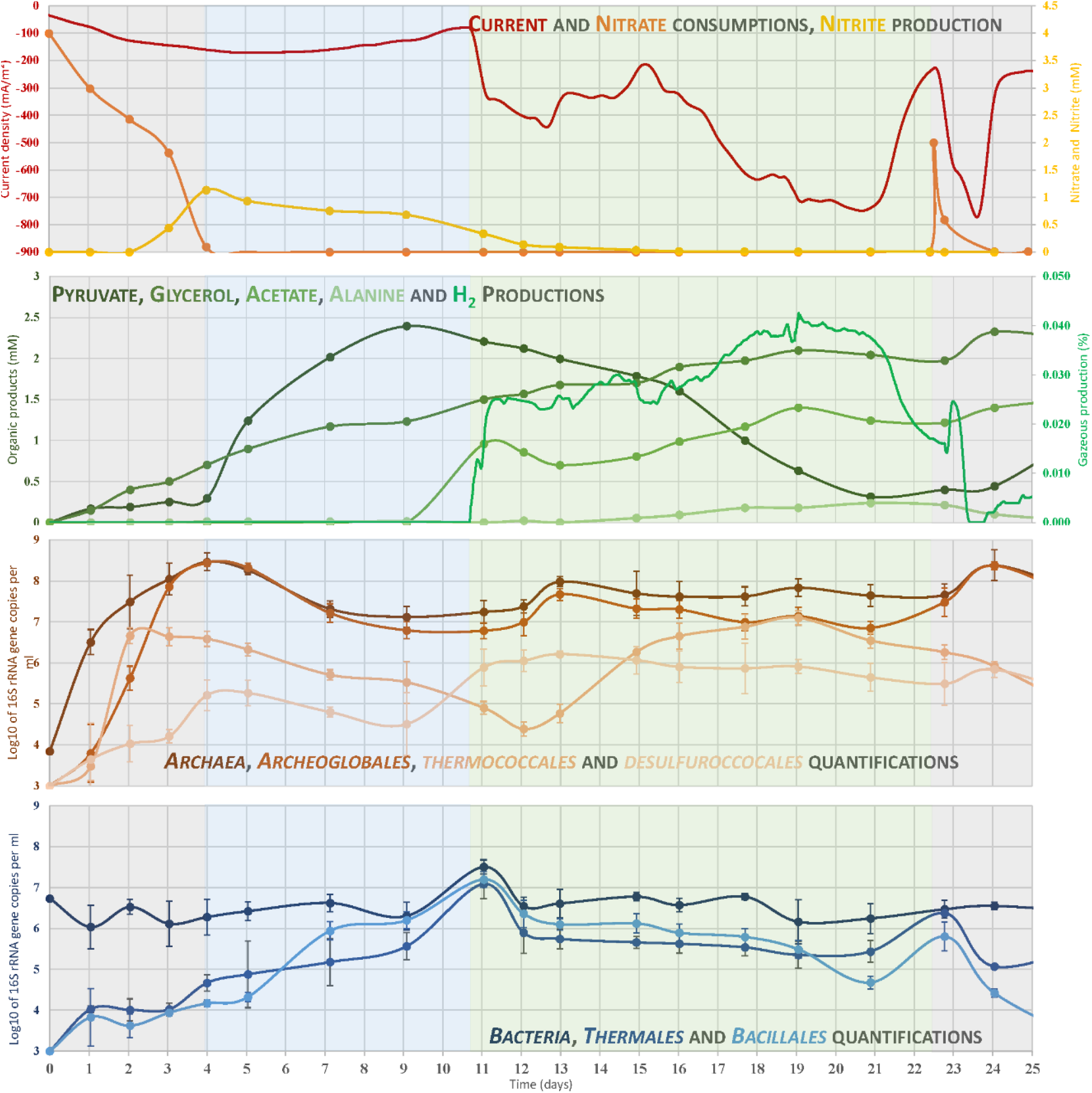
Consumption, production, and microbial 16S genes quantification over time of the enrichment of Nitrate-reducing eletrotrophic community in Microbial Electrochemical System at 80°C inoculated with 0.5% (w/v) of a crushed chimney sample. Fig. 1A: current, nitrate and nitrite consumptions; Fig. 1B: product productions; Fig. 1C: qPCR evolution from Archaea specific primers; Fig. 1D: qPCR evolution from Bacteria specific primers.

### Nitrate/Nitrite quantification

Nitrate consumption was analyzed using a wet oxidation technique with automated colorimetry as described in [22] after culture medium centrifugation at 14000 rpm for 5 min.

### Identification and quantification of organic compounds production

To identify and quantify organics productions from the biofilm, samples of liquid media were collected every 24 to 48h and analyzed by ^1^H NMR Spectroscopy. Four hundred microlitres of each culture medium were added to 200 μl of PBS solution prepared in D_2_O (NaCl, 140 mM; KCl, 2.7 mM; KH_2_PO4, 1.5 mM; Na_2_HPO4, 8.1 mM, pH 7.4) supplemented with 0.5 mM of trimethylsilylpropionic acid-d4 (TSP) as NMR reference. All the 1D ^1^H NMR experiments were carried out at 300 K on a Bruker Avance spectrometer operating at 600 MHz for the ^1^H frequency and equipped with a 5-mm BBFO probe.

Spectra were recorded using the 1D nuclear Overhauser effect spectroscopy pulse sequence (Trd-90°-t1-90°-tm-90°-Taq) with a relaxation delay (Trd) of 12.5 s, a mixing time (tm) of 100 ms and a t1 of 4 μs. The sequence enables an optimal suppression of the water signal that dominates the spectrum. One hundred twenty-eight free induction decays (FID) of 65,536 data points were collected using a spectral width of 12 kHz and an acquisition time of 2.72 s. For all the spectra, the FIDs were multiplied by an exponential weighting function corresponding to a line broadening of 0.3 Hz and zero-filled before Fourier transformation. NMR spectra were manually phased using Topspin 3.5 software (Bruker Biospin Corporation) and automatically baseline corrected and referenced to the TSP signal (δ = −0.015 ppm) using Chenomx NMR suite (version 7.5) software (Chenomx Inc.). A 0.3 Hz line-broadening apodization was applied prior to spectral analysis and ^1^H-^1^H TOCSY [23] and ^1^H-^13^C HSQC [24] experiments were recorded on selected samples in order to identify the detected metabolites. Quantification of identified metabolites was done using Chenomx NMR suite (version 7.5) software (Chenomx Inc.) using the TSP signal as an internal standard.

### Biodiversity analysis

The taxonomic affiliation was performed according to [25]. DNA was extracted from 1 g of the crushed chimney and at the end of each period of culture from the scrapings of half of the working electrode and from centrifuged pellets of 50 mL of spent media. The DNA extraction was carried out using the MoBio PowerSoil DNA isolation kit (Carlsbad, CA). The V4 region of the 16S rRNA gene was amplified using the universal primers 515F (5′-GTG CCA GCM GCC GCG GTA A-3′) and 806R (5′-GGA CTA CNN GGG TAT CTA AT-3′) [26] with Taq&Load MasterMix (Promega). PCR reactions, amplicons sequencing, and taxonomic affiliation were carried as previously described [21]. To analyze alpha diversity, the OTU tables were rarefied to a sampling depth of 13190 sequences per library, and three metrics were calculated: the richness component (number of OTUs), the Pielou’s index and Shannon’s biodiversity index. Rarefaction curves are presented in Supporting Information S2. The raw sequences are available on the European Nucleotide Archive (accession number: PRJEB35427).

### Microscopy observation with fluorescent in-situ hybridization (FISH)

Prior performing microscopic observation (Fig. 3), the working electrode from the end of each experiment was fixed with 2% paraformaldehyde and kept at 4 °C. To highlight the presence and abundance of microbes from different domain and order, fluorescently labelled (CY3, FITC) probes (Biomers.net GmbH, Germany) were used to label nucleic acids (Syto9), *Bacteria* (EUB338-FITC), *Archaea* (ARCH917-CY3), *Euryarchaeota* (Eury806), *Crenarcheota* (Cren537), *Thermococcales* (Tcoc164) and *Archeoglobales* (Arglo32) [27, 28]. Samples of working electrode or cells on the filter were incubated with fluorescent probes in an equilibrated humidity chamber at 48°C for 2–6 h and then wash with washing buffer at 42 °C for 15min. Samples were then dried in 80% ethanol and mount on a glass slide with antifadent AF1 (Citifluor, USA) added with 4’-6’-diamidino-2-phenylindole (DAPI) (Sigma Aldrich) for counterstain at a final concentration of 2 μg.ml^−1^. Samples observation were performed on a confocal LSM780 microscope (ZEISS, Germany) equipped with an x10, EC PLAN-NEOFLUAR objective. Blue, green and red fluorescence emissions were acquired by excitation at 358, 488 and 561 nm, respectively, using three lasers. Image stacks (at 0.5–1 μm steps) were acquired with GaAsP photomultiplier tube detectors. Epifluorescence micrographs were processed using the Zen software (Zeiss, Germany).

### Quantitative PCR of phylogenetic orders

Quantification of *Bacteria*, *Archaea* and specific phylogenetic orders retrieved in MiSeq analysis were carried out by qPCR method. SsoAdvanced™ Sybr Green Supermix was used on a CFX96 Real-Time System (C1000 Thermal Cycler, Bio-Rad Laboratories, CA) with the primers specified in table 1. 16S rRNA primers were specifically and manually designed for quantification of most represented orders using MEGA6 Software [29] and 16S rRNA reference sequences from Silva database (http://www.arb-silva.de). Newly designed primers were validated *in silico* on TestPrime software (https://www.arb-silva.de) and on different cultures of species belonging or not to targeted order. The PCR program was composed of a 5min initial denaturation step at 98°C followed by 50 cycles of 10s denaturation step at 98°C, a hybridization step of 20 s at temperature indicated in table 1 and a 40 s elongation step at 72°C, with melting curves generated at the end of each reaction to ensure product specificity. A standard curve from 10^2^ to 10^10^16S rRNA gene copies was obtained by diluting pGEM-T plasmids harboring 16 rRNA gene fragments specific to bacteria, archaea or specific members of each order. qPCR quantifications were compared to quantification on 1 ml culture of targeted species filtered on a 0.2 μm filter and stained with DAPI. Average cell count per ten optical fields has been reported to the total area of the filtration zone and then to 1ml of culture. Results of qPCR were expressed in a number of 16s rRNA gene copies per milliliter of liquid media. Due to the qPCR protocol, a minimum threshold of detection of 3 log_10_/ml was observed.

**Table 1:**
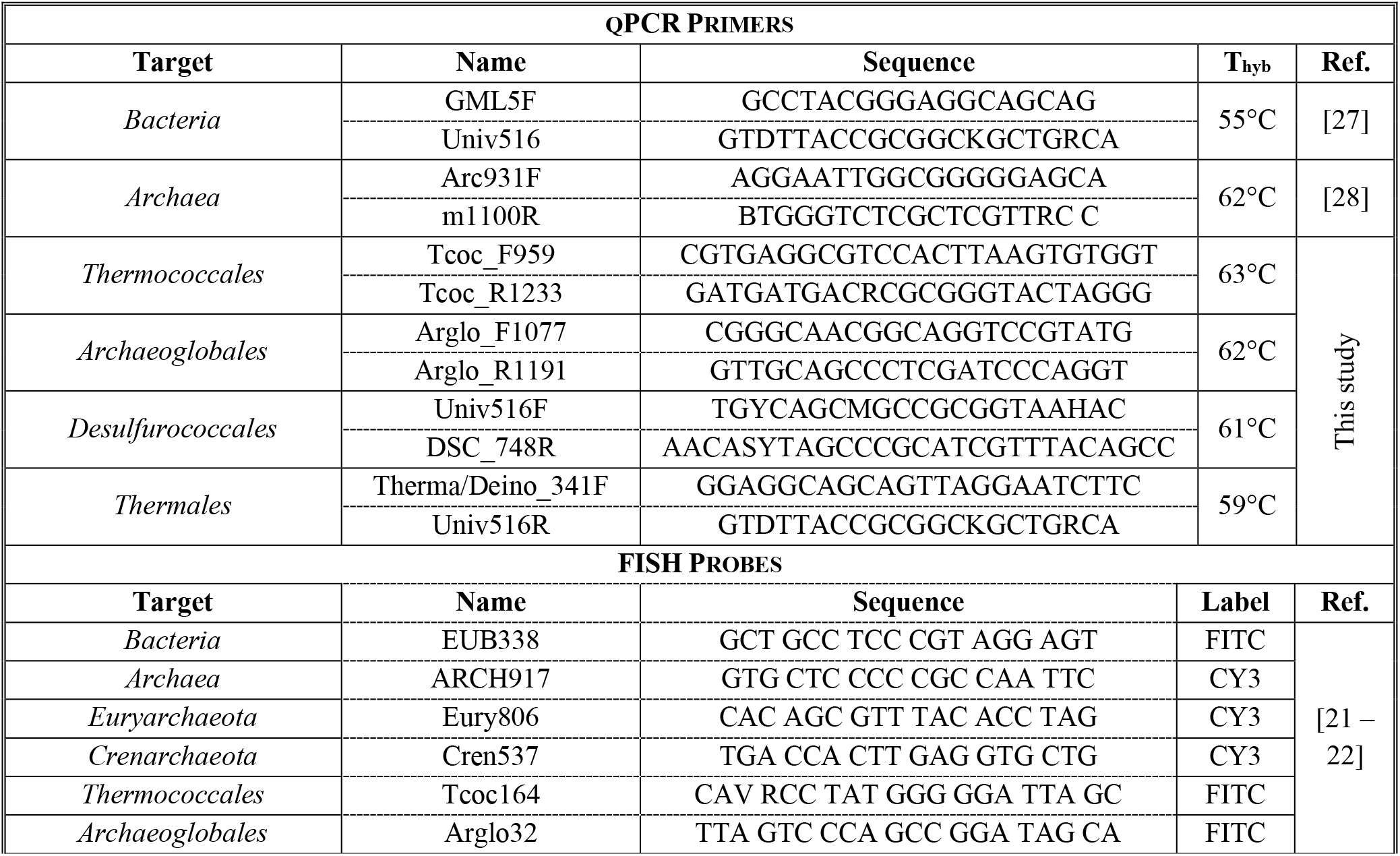
16S rRNA primer for qPCR quantification of Bacteria, Archaea, specific phylogenetic orders and 16S rRNA labeled probes for FISH Microscopy

## Results

### Current and Nitrate consumptions

A preliminary characterization of our system was carried out to control the growth conditions of our electroautotrophic biofilm. First, the organic matter present in our inoculum was measured by NMR, showing less than 75 μM cumulated of potential electron donors and acceptors. Then, an open circuit potential (OCP) control was performed in the same conditions as our enrichment and inoculated with the hydrothermal chimney. In this control, qPCR, NMR and microscopy measurements didn’t show any growth over time (data not shown). Thereby, in the MES filled with the mineral medium, the cathode poised at −590 mV vs SHE was the only potential energy source available for microbial growth. Under this experimental condition, CO_2_, continuously sparged into the reactor, was the only available carbon source and NO_3_, present in mineral medium, was the only electrons acceptor source available. Evolution of nitrate, nitrite and current consumption are reported Fig. 1A. The nitrate reduction can be divided into two phases. During the first four days, the nitrate is consumed (from 4 mM to 0.22 mM) correlated with low production of nitrite (up to 1mM). After D4 (day 4^th^), a slow consumption of nitrite (from 1mM to 0.33mM) was observed until the D12. During these first two phases, a slow increase of current consumption was measured (up to −180 mA.m^−2^) while nitrate and nitrite are consumed. Then, a drastic increase in current consumption was observed to reach −750 mA/m^2^ until D20. At D22, the addition of 2 mM of nitrate, which is rapidly reduced, has allowed to reach back the maximum current consumption (−765 mA/m^2^) before a quick drop again.

### Electrosynthesis of organic compounds

During the growth of the nitrate-reducing electrotrophic community, different organic compounds were released in the liquid media (Fig. 1B) while none were detected in the controls. Analysis of liquid samples over time by RMN method allowed to identify and quantify the production of a significant amount of acetate, glycerol, pyruvate, and alanine. Glycerol was continuously and slowly produced over time to reach 2.3 mM at D24. Pyruvate started to be strongly produced at D4, to reach a maximum of 2.4 mM at D9 followed by a decrease (until 0.25 mM; D21) due to a microbial consumption over time. This pyruvate production occurred following the total depletion of nitrate in the liquid media. Acetate started to be produced at D11 to reach a maximum (1.4 mM; D19) correlated with the pyruvate decrease, the current consumption, and H_2_ production. Moreover, low production of alanine was detected after D11 to reach a maximum (0.12 mM; D21) and decrease later.

H_2_ production (Fig. 1B) has appeared strongly related to the variation of current consumption (Fig. 1A). This production has increased quickly in 24h up to 0.025 ml.min-1, remained stable for five days and then slowly increased to reach a maximum of 0.04 ml.min-1 at D19. This production decreased from D21, correlated with the depletion of the pyruvate leading to the decrease of current consumption (−220 mA/m^2^; D22).

Others organic compounds have been produced during this experiment. Peaks of methanol and ethanol were measured, that reached a maximum of 9 mM and 0.16 mM, respectively at D4 (data not shown). Moreover, slow production of acetamide (maximum of 69 μM at D12), benzoate like molecule (slow increase up to 0.13 mM at D21), 2-aminoisobutyric acid (maximum of 22 μM at D12) and formate (variations between 8 to 70 μM between D1 and D4) were also observed over the experimentation period (data not shown).

These H_2_ and organic compounds productions were not observed in abiotic and OCP controls with nitrate, meaning that this production is catalyzed by the growth of microorganisms on the electrode.

### Enrichment over time of dominant phylogenetic orders

To study the evolution of the composition of the electrotrophic community over time (Fig. 1C & 1D), a quantification by qPCR was performed with primers specific to each significant order (Table 1). The quantifications were performed on harvested liquid samples over time. Due to the presence of only the electrode as an energy source, microorganisms found in liquid media necessarily arise from on-electrode biofilm release. During the first four days, a quick increase of archaeal 16S rRNA gene copies (Fig. 1C) was observed (from 3.8 to 8.4 log_10_), simultaneously to the increase of *Archaeoglobales* for four days (from <3 to 8.4 log_10_) and *Thermococcales* for two days (from <3 to 6.6 log_10_). These increases corresponded to nitrate reduction and nitrite accumulation (Fig. 1). Between D3 and D4 *Desulfurococcales* 16-rRNA-gene copies have increased correlated to the acceleration of the nitrate reduction and nitrite production. No increase in bacterial 16S-rRNA-gene copies was observed during this period. From D3 to D11, *Archeoglobales, Desulfurococcales*, and *Thermococcales* 16S-rRNA-gene copies slowly decreased of 1-1.5 log_10_ at the same time as the pyruvate accumulation in the liquid media. During this period, the *Thermales* 16S-rRNA-gene copies increased (from 4 to 7 log_10_) related to the nitrite reduction (Fig. 1A & 1D). From D11 to D22, *Thermococcales* and *Desulfurococcales* species were enriched while other orders slowly decreased or maintained in the liquid media. The enrichment of the latter was related to the decreasing of pyruvate and production of acetate, hydrogen and alanine (fermentation products; Fig. 1B). It is worth noting that *Archaeoglobales* species increased also between D11 and D13 correlated to the weak degradation of acetate (from 1 mM to 0.6 mM). Finally, during the last days, the nitrate addition allowed the enrichment of *Archeoglobales* species followed by heterotrophic *Thermales*.

### Statistical analysis of the correlation between variables

A Principal Component Analysis (PCA) was performed on to study the correlation between the current density, the nitrate-, nitrite- and organics concentrations as well as microbial communities (Fig. 2). It represents the distribution of each sample (represented by the day number post-inoculation) and the contribution of each variable on a Biplot composed of the two first dimensions, explaining respectively 39.9% and 18.6 % of total variances. These two dimensions allowed to discriminate four temporal groups, called phases in this study. The first phase was represented by samples from D0 to D4 of culture, mainly explained by the *Archaeoglobales* evolution and the nitrate consumption but also inversely correlated to the pyruvate and fermentation products (Fig. 2). The second phase between D5 to D11, is mainly explained by the evolution of *Thermales*, nitrite, and accumulation of pyruvate. The third phase from D12 to D22, mainly explained by the first dimension, was linked to the accumulation of fermentation products (alanine, acetate, H_2_) and current consumption. Finally, the fourth and last group, represented by the samples of D23 and D24, was significantly different from other groups but didn’t link specifically to some parameters. It is worth noting that the contribution of *Thermococcales* and *Desulfurococcales* was very weak in the building of the 2 dimensions of the PCA. Indeed, their evolutions could not be linked to one temporal group in the PCA analysis. This can be explained by their evolution in more than one phase or independently of other variables.

**Figure 2:**
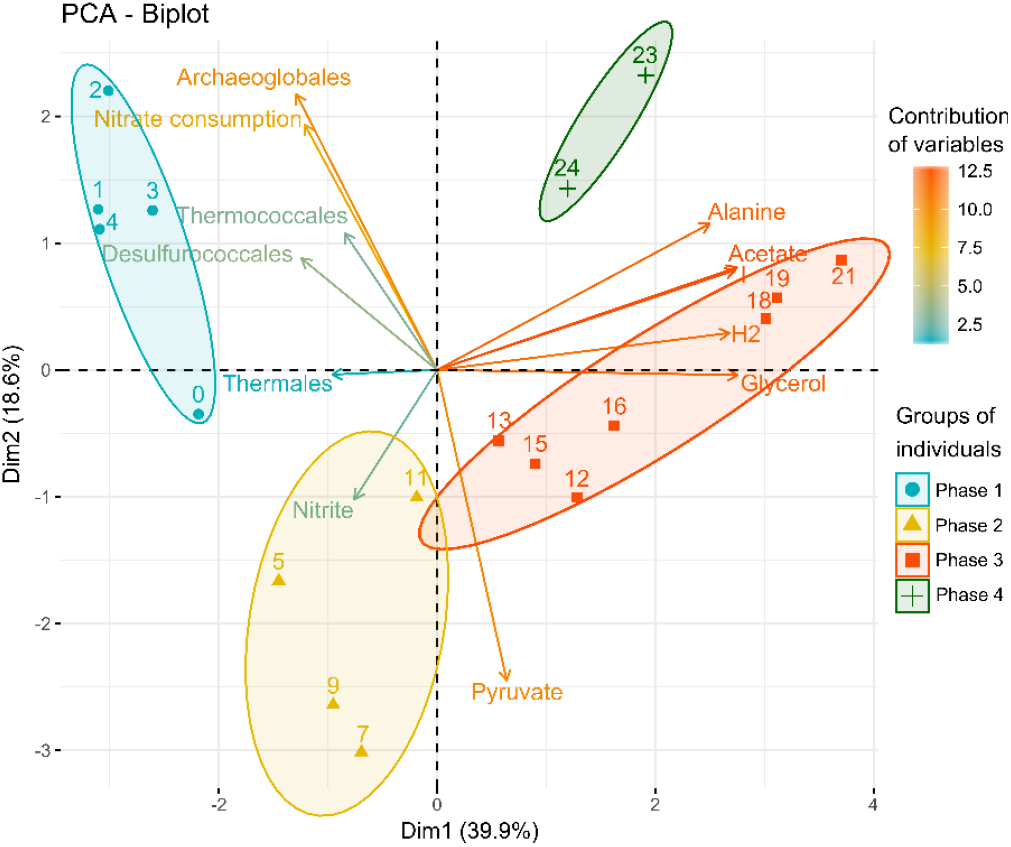
PCA Analysis of nitrate, nitrite, organic measurements, and evolution of quantity (qPCR) of dominant phylogenetic orders over the days of culture. It represents the distribution of each sample (represented by the day number post-inoculation) and the contribution of each variable on a Biplot composed of the two first dimensions, explaining respectively 39.9% and 18.6 % of total variances. These two dimensions allowed to descriminate four temporal groups.

**Figure 3:**
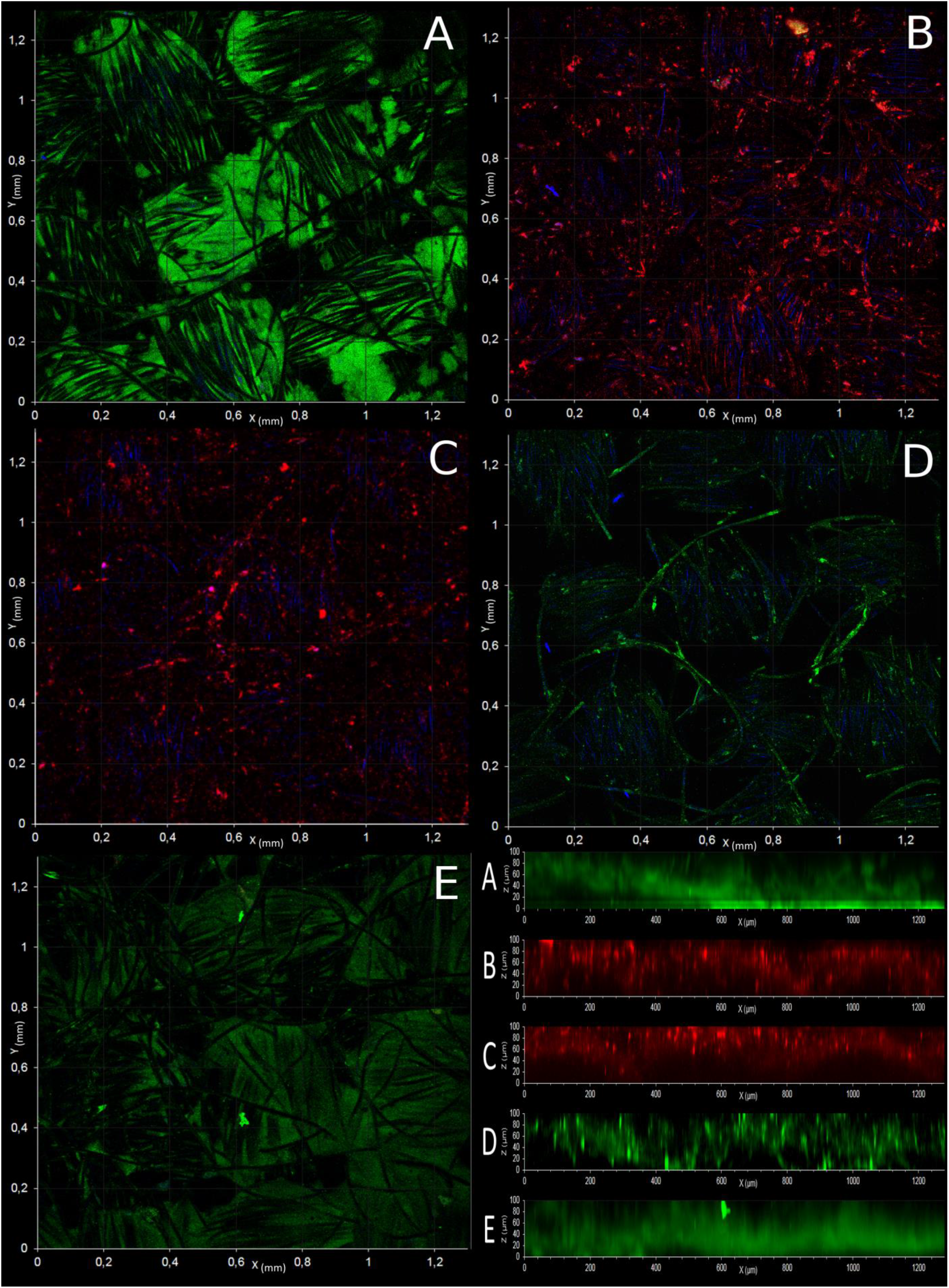
Confocal observation and depth profile (F) of mature microbial biofilm developed on cathode, stained with FISH probes (Table 1) specific to Bacteria (A, green), *Euryarchaeota* (B, red), *Crenarchaeota* (C, red), *Thermococcales* (D, green) and *Archeoglobales* (E, green) at Obj. x100. Blue signal corresponds to the reflectance of carbon fiber.

### Microscopic observation of electrotrophic biofilm

Confocal observation of electrotrophic biofilm on the electrode of the first experiment was performed with FISH probes specific to *Bacteria*, *Archaea*, *Euryarchaeota*, *Crenarchaeota*, *Thermococcales* and *Archeoglobales* (Fig 3). It allowed highlighting the preferential presence of *Bacteria* (*Thermales*) and *Archeoglobales* in the depth of electrode (Fig 3A, 3E and 3F), while *Thermococcales* and *Crenarchaeota* (*Desulfurococcales*) (Fig 3C and D) are growing mostly on the external surface of the electrode. These observations suggest the colonization of nitrate-reducing microorganisms (e.g. some *Archaeoglobales* and *Thermales*) in the depth of the electrode while fermentative microbes (e.g. *Thermococcus* and *Thermodiscus* sp.) growth in peripheral of this colonization.

### Biodiversity on electrode and liquid media over experiments

Once a first colonization of the electrode was obtained, we aimed to study if this community could lead to the development of a more complex and mature ecosystem over time, as observed in the older hydrothermal chimneys. Thus, two successive subcultures (cf. Material and methods) were performed in new MES inoculated with media from the previous enrichment. After stabilization of current consumption, the communities present on electrodes (C1-, C2- and C3-WE) and in liquid media (C1-, C2- and C3-LM), were identified through 16S Metabarcoding (Fig. 4). While the environmental sample showed large biodiversity with a good relative evenness (Shannon index = 5.19 and Pielou’s evenness index = 0.695), the three enrichments have shown an increase of the enrichment of microorganisms mainly in the liquid media (Shannon indexes: C1-LM=1.92; C2- LM=4.16, C3-LM=5.04). The relative evenness in these enrichments showed an equal good distribution of species (evenness indexes: C1-LM= 0.316; C2-LM=0.618; C3-LM=0.752). However, the enrichment on electrode remained equivalent over the subcultures (Shannon index ~3.3 and evenness index ~0.49).

**Figure 4:**
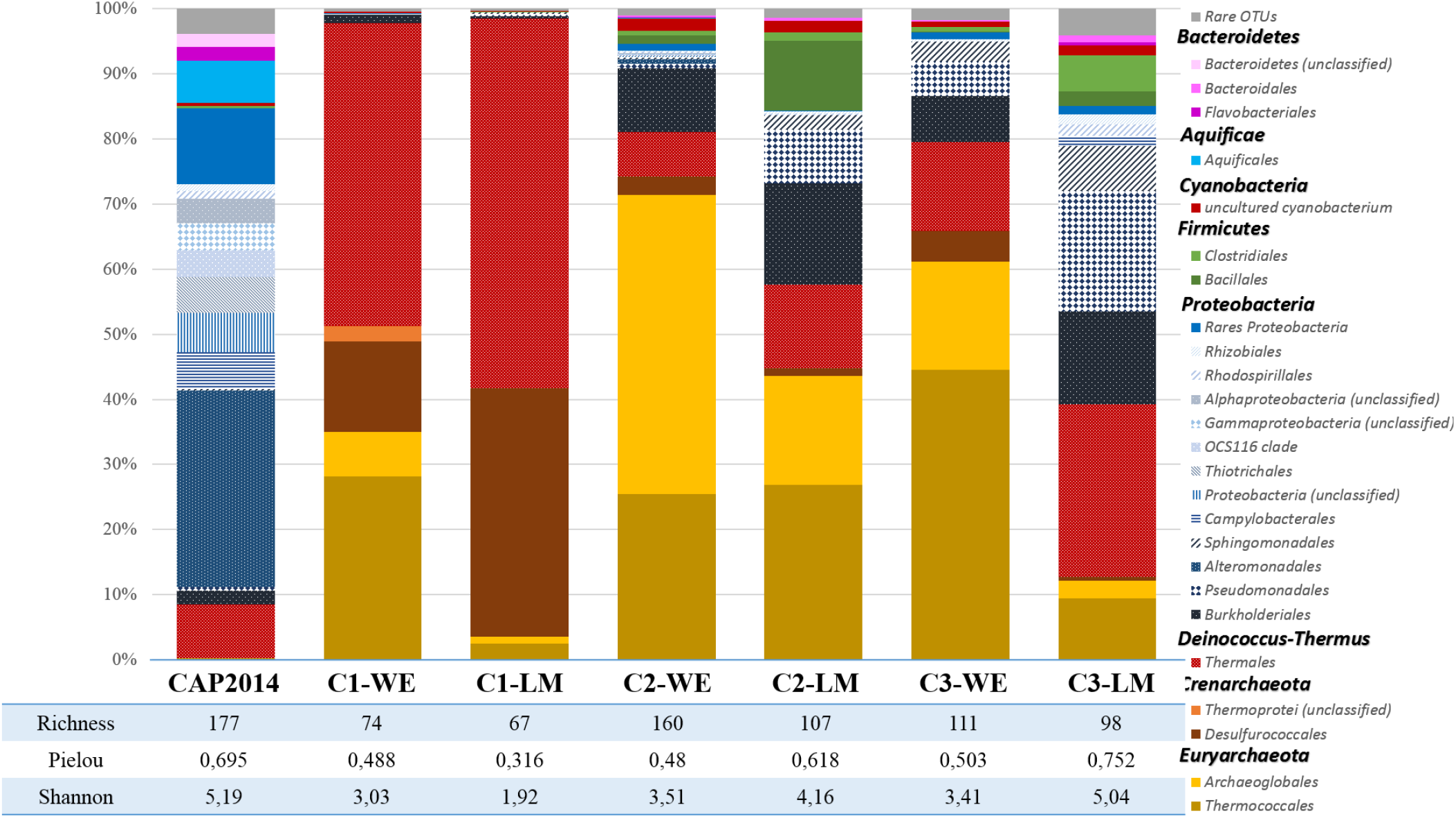
Dominant taxonomic affiliation at the order level and biodiversity indices of enriched microbial communities from a crushed chimney sample, as plotted on the cathodic working electrode (WE) and liquid media (LM) over the subcultures. OTUs representing less than 1% of total sequences of the samples are pooled as ‘Rare OTUs’.

Based on average abundance analysis (Fig. 4), the microbial diversity in the first culture (C1-WE and C1-LM) was mainly dominated by species belonging to 4 orders: *Thermococcales* (28.2 – 2.5 %), *Archeoglobales* (6.8 – 1 %), *Desulfurococcales* (13.9 - 38.1 %) and *Thermales* (46.5 – 56.8 %) respectively on electrode and in liquid media. The remaining biodiversity represents only 0.46 % on the electrode and 0.16% in liquid media, shared between *Proteobacteria*, *Firmicutes* and *Actinobacteria* species. The second culture (C2-WE and C2-LM) showed specific enrichment of *Archeoglobales* on the electrode (45.8%) and in liquid media (16.8%). Moreover, the *Thermococcales* remained stable on the electrode (25.5%) while it enriched in liquid media (26.8%). By contrast, the proportions of *Desulfurococcales* (2.8 – 1.2%) and *Thermales* (6.85% - 12.7%) have drastically dropped on the electrode and liquid media. Interestingly, we observed an increase of the remaining biodiversity with enrichment of *Burkholderiales* (9.75% - 15.64%), *Pseudomonadales* (8.22% only in liquid) and *Bacillales* (10.7% only in liquid) in this second subculture. Finally, the last enrichment (C3-WE and C3-LM) showed the massive growth of *Bacteria* in liquid media (87.3%), mainly affiliated to *Thermales* (26.4%), *Burkholderiales* (14.3%), *Pseudomonadales* (18.4%), *Sphingomonadales* (6.9%) and *Clostridiales* (5.46%) while the composition of the biofilm on electrode is substantially the same as in the second enrichment. The taxonomic affiliation of dominant OTUs has allowed identifying the *Thermococcales* as closely related to around 20 *Thermococcus* spp., *Archeoglobales* to *Geoglobus ahangari* strain 234 (99% similarity), *Archaeoglobus sp.* Fe70 (99% similarity), and *Ferroglobus placidus* (98% similarity) and *Desulfurococcales* to *Aeropyrum* sp. AF1T6.18 (98% similarity) and *Thermodiscus maritimus* (97% similarity). For *Bacteria*, *Thermales* are composed of 2 OTUs, mainly affiliated to *Vulcanithermus mediatlanticus* strain TR (99% similarity) and a new species of *Thermaceae* (close at 95% to *Vulcanithermus* sp. BF2T511) respectively, *Bacillales* to *Geobacillus thermodenitrificans*, *Burkholderiales* to *Ralstonia* spp. and *Pseudomonadales* to *Pseudomonas* spp.

## Discussion

The monitoring in our conditions of the current consumption, the decrease and transformation of the electron acceptor, the production of organic intermediates and the evolution of the community are proofs of the development of a hyperthermophilic electrolithoautotrophic community on electrode. To identify the first colonizers of this community and the metabolisms involved, an analysis of the correlations between the different parameters was performed. The PCA analysis (Fig. 2) and the observations of the evolution of the parameters (Fig. 1) allowed to separate the experiment in 4 phases

### Phase 1: Electrotrophic hyperthermophilic nitrate reduction

The first phase, from inoculation to D4 (Fig. 1A), corresponds to the nitrate reduction with a transient nitrite accumulation while current is consumed. This is related to the growth of *Archaeal* species, mainly *Archaeoglobales* immediately followed and surpassed by *Thermococcales* and then *Desulfurococcales* species (Fig. 1C). Alongside, low production of organics is observed, mainly glycerol and pyruvate and a weak production of H_2_. Some species of the order *Archaeoglobales* (*Geoglobus ahangari* and *Ferroglobus placidus*) have already been described to perform extracellular electron transfers by using an electrode as electron acceptor [30]. How Archaea carry out exogenous electron transfer is still unknown [31] but the electrotrophic metabolism is recognized when a current consumption is observed [32, 33]. Among the three enriched orders, *Archaeoglobales* are the first species growing from CO_2_ as carbon source and electrode as the electron donor. The taxonomic affiliations (Fig. 4; C1-WE-LM) of two dominant *Archaeoglobales* OTUs obtained after 24 days of growth are closely related to the three genera belonging to these orders: *Geoglobus* spp., *Archaeoglobus* spp. and *Ferroglobus* spp. These three genera are known to grow lithoautotrophically through the reductive acetyl-CoA/Wood-Ljundahl pathway. However, *F. placidus* is currently the only member of the *Archaeoglobales* which has been shown to be able to use nitrate as electron acceptor accompanied by transient NO_2_ ^−^ accumulation and production of N_2_O [34]. Nonetheless, a cluster of genes encoding a putative nitrate reductase has though been identified in *Geoglobus ahangari* and different *Archaeoglobus* spp. [35–37] but the physiological evidence of nitrate reduction is still missing in conditions of classical cultures. We can suggest here, that *Archaeoglobales* (*Geoglobus* spp., *Archaeoglobus* spp. or *Ferroglobus* spp.) grow on electrode and use the nitrate as an electron acceptor with cathode as reducing power allowing the production of biomass and the release of organic compounds from CO_2_ reduction during the first four days. These compounds could serve as a source of carbon and energy for different heterotrophic microorganisms.

Alongside, *Thermococcales*, whose OTUs were affiliated to 38 validated species of *Thermococcus*, increased during the first three days (Fig. 1C). All members of the *Thermococcales* are characterized by their ability to use complex or simple peptides as energy and carbon source by necessarily using elemental sulfur as electron acceptor [38]. The breakdown of peptides and amino acids leads to the subsequent production of organic acids linked to substrate-level phosphorylation. Many *Thermococcales* species can also grow by fermentation of various carbohydrates without the need for S° [39]. The major fermentation products are acetate, H_2,_ and CO_2_. However, the organic acid such as acetate didn’t accumulate despite the significant growth of *Thermococcales* (7.9 log) in the liquid medium (D1 to D4). Moreover, representatives of *Thermococcales* have never been shown to be able to reduce nitrate. Notwithstanding, some *Thermococcales* have been described as carboxytrophic which use the oxidation of carbon monoxide (CO) to produce CO_2_ and H_2_ [40, 41]. The carbonyl branch of the reductive acetyl-CoA pathway, present in *Archaeoglobales*, is known to start from the reduction of CO_2_ to CO by a bifunctional CO dehydrogenase (CODH). In excess of electron donor or CO_2_, CO can be released in liquid media [42]. In microbial electrosynthesis system, overfeeding of electrons from the polarized cathode (here at −590 mV vs ESH) might favor the production and release of CO by *Archaeoglobales* on the surface of the electrode, that diffuses in the liquid medium afterwards. The transient and weak accumulation of H_2_ could be explained by this metabolism described for several *Thermococcus* [40, 41]. However, neither autotrophic growth has not been demonstrated nor have autotrophic pathways been predicted in the available genome sequences. Indeed, carboxydotrophic growth of *Thermococcales* requires a source of organic carbon in the form of peptides [43, 44] but we assume that the growth of *Archaeoglobales* could supply the organic carbon source through the excretion of extracellular polymeric substances that would promote *Thermococcales* growth.

Alongside with *Archaeoglobales* and *Thermococcales* growths and nitrate consumption, the growth of heterotrophic *Desulfurococcales* and *Thermales* have been observed (Fig. 1C & 1D). While only one species of *Desulfurococcales*, *Pyrolobus fumarri*, have been shown to reduce nitrate [45], most of *Thermales*, such as *Vulcanithermus* spp, or *Oceanithermus* spp. are able to reduce nitrate and even reduce nitrite for some [46]. The delay before *Desulfurococcales* and *Thermales* emergence suggest their heterotrophic growth from syntrophic relations. These trophic relations would have been set by fermentation of organics produced and/or by partially performing the last step of the dissimilatory nitrate reduction from nitrite produced by nitrate-reducing microorganisms.

### Phase 2: Electrosynthesis of pyruvate

The second phase is marked by the increase of pyruvate production (Fig. 1B). This effect only occurred significantly from D4, while the nitrate reached its complete depletion and *Archeoglobales* stop growing. Thus, we can suggest that pyruvate production is attributed to lithoautotrophic *Archeoglobales* fixed on the electrode, closely affiliated to *Geoglobus* spp., *Ferroglobus* spp. and *Archaeoglobus* spp. As previously mentioned, these *Archaeoglobales* use the Wood Ljungdahl pathway for carbon fixation through the bifunctional CODH/acetyl-CoA synthase complex, which allows to link methyl and carbonyl branches and enable acetyl-CoA production [47]. Acetyl-CoA can be used directly for biosynthesis or converted to pyruvate by the reversible reaction of pyruvate:ferredoxin oxidoreductase (PFOR) EC: 1.2.7.1] [48]. This conversion of CO_2_ to pyruvate uses reduced ferredoxins, normally regenerated by hydrogenases [34, 49]. Then, this pyruvate is normally used for biosynthesis through conversion to oxaloacetate by pyruvate carboxylase [50] which uses one mole of ATP. But, in absence of electron acceptor, the excess of electrons continuously injected into the cells through the cathode (−590 mV vs ESH) lead to the accumulation of reduced electronic shuttles and to the production of pyruvate (Fig. 5). This continuous production of pyruvate is however not enough energetic to allow the growth of the *Archaeoglobales* [49] and no ATP is then available for the conversion to oxaloacetate. However, *Archaeoglobales* seems to sustain during several days (D4 to D11) with a new type of energetic coupling that remains to be fully explored. Interestingly, *Thermococcales* have stopped to grow (D2 to D11) while they are known to be able to perform pyruvate fermentation to acetate through the oxidative decarboxylation of the pyruvate by the pyruvate:ferredoxin oxidoreductase (POR) 39, 40]. The transient accumulation of nitrite could cause the inhibition of POR preventing the pyruvate fermentation as already observed in *Pyrococcus furiosus* [51].

**Figure 5:**
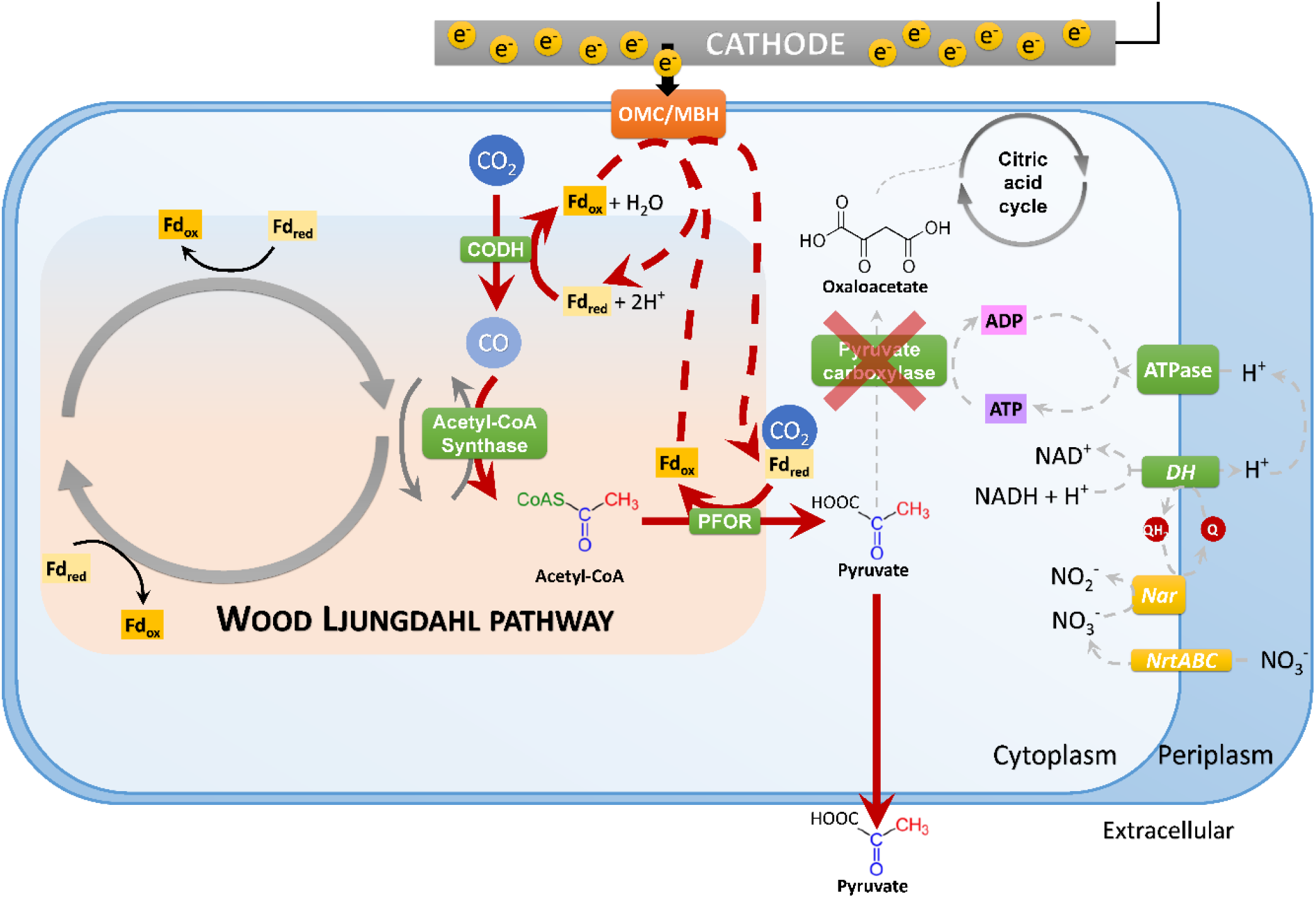
Electrosynthesis pathway through the Wood-Ljungdahl carbon fixation pathway. Red dot lines represent the hypothesis of the inhibition of those reactions in the absence of electron acceptor and a overfeeding of electrons (conductive chimney walls or polarized cathode).

### Phase 3: Electrofermentation of pyruvate

The third phase of our experiment (D11 to D22.5) is characterized by an increase of current consumption and the fermentation of pyruvate into acetate, alanine, and H_2_ (Fig. 1A & 1B). During these phenomena, *Thermodiscus* and *Thermococcus* populations increase in liquid media. *Thermodiscus* species are poorly studied but are described as obligatory heterotrophic *Crenarchaeota* able of sulfur respiration and fermentation on complex organic compounds [52]. *Thermococcales* are also known to be heterotrophic and fermentative *Euryarchaeota*. Nitrite is known to inhibit fermentation in Thermococcales [51]. Thus, *Thermococcus* and *Thermodiscus* start to grow through pyruvate fermentation only after nitrite is depleted in D11, explaining the previous accumulation of pyruvate during the inhibition. Interestingly, this fermentation from the D11 is related to current consumption, and to the growth of *Archaeoglobales* despite the electron acceptors lack. *Thermococcales* and *Thermodiscus* are believed to not be able to grow on cathode as they cannot use electron acceptors and cannot fix CO_2_. Thus, this current consumption could be attributed to the mechanism of Direct Interspecies Electron Transfers (DIET) already observed in various co-cultures [54, 55]. The FISH microscopic observations (Fig. 3) have shown the peripheral development of *Thermococcales* and *Desulfurococcales* on the electrode around *Archaeoglobales*, which were in close contact with the carbon fibers. This spatial configuration would have allowed direct contacts between the electrode, *Archaeoglobales*, and fermentative microorganisms. In this hypothesis, *Thermococcales* would have served as an electron acceptor for *Archaeoglobales*, allowing them to grow again in the absence of nitrate while supplying *Thermococcales* with electrons for their metabolism. The electron flow introduced by *Archaeoglobales* to *Thermococcales* would have led to the electrofermentation of pyruvate linked to current consumption and the production of acetate, alanine, and H_2_ [56]. Thereafter, these compounds could be used as electrons donors for *Archaeoglobales* [57], in addition to the electrode’s electron-downflow. Thus, the established DIET would have allowed the concomitant growth of *Archaeoglobales* and *Thermococcales* through electron transfer (D11 to D13). This suggests that the *Archaeoglobales* and *Thermococcales* would be much more metabolically diverse in the hydrothermal vent systems with various acceptors or potential electron donors.

### Phase 4: The trophic chain from electro-autotrophic nitrate reduction to heterotrophy

In the 4^th^ and last phase, the successive addition of nitrate after D22 (Fig. 1), showed the concomitant course of the different metabolisms with consumption of nitrate, an increase of heterotrophic then autotrophic nitrate-reducing microorganisms, alongside with the production of fermentation products (H_2_ and acetate). This indicates that in mature communities, metabolic plasticity is developed to allow trophic interaction between auto- and heterotrophs but also respiring (including DIET) and fermentative microorganisms to allow their mutual survival in harsh and quickly changing conditions in hydrothermal context.

### Thereafter: Development of more complex ecosystem from deep hydrothermal vents

After the 4^th^ phase, the electrode and the mineral medium of MES have been renewed then inoculated to 1% (v/v) with the harvested liquid media during the 4^th^ phase. The current density has been monitored during one week in the same previous conditions (nitrate, CO_2_ and polarized electrode to −590mV vs. SHE), then the electrode and liquid media have been harvested to analyze the enriched biodiversity. Another similar culture was performed in the same conditions, by using the harvested liquid media of the previous culture. During all these cultures, the electrode was the sole energy source, with only less than 0.3 mM of cumulated organic products (acetate, glycerol and pyruvate) and 7μg of estimated biomass (qPCR x mass of a cell) brought by the inoculum from the previous enrichment. The MiSeq results exhibited the enrichment of a part of the biodiversity initially present in the inoculum (Fig. 4). Subcultures of electrotrophic biofilm from successive planktonic cells allow the development over time of more and more diverse species in liquid media and on the cathode. These species growing late are affiliated to *Proteobacteria*, mainly members of *Burkholderiales*, *Pseudomonadales*, and *Sphingomonadales*. These bacterial orders are only represented by heterotrophic species. Thus, their late development after the autotrophic condition of the experiment suggests trophic interaction between electro-autotrophic and heterotrophic species.

## Conclusion

As supposed by Yamamoto et al. 2017, electricity generation in deep hydrothermal systems is expected to affect surrounding biogeochemical processes and the development of microbial communities. Here, we have demonstrated that some microorganisms are able to use the available electrons on the surface of polarized electrode as an alternative energy source to other and conventional electron donor molecules, such as molecular hydrogen, hydrogen sulfide or methane. Thus, some microbial populations, particularly those belonging to the *Archaea* domain, might grow from the electrons produced by the spontaneous and generalized production of electricity between the hydrothermal fluids and the seawater through the hydrothermal chimneys. The presence of these specific microorganisms, that can be described as electrolithoautotrophic microorganisms, highlights a new trophic type considered as electrotrophic compared to the chemotrophic and phototrophic ecosystems found on our planet. In addition, the probable presence of DIET within the studied microbial populations, allows extrapolation of these results on a metabolic alternation in these ecosystems, which would lead the electroactive species of *Archaeoglobales*, *Thermococcales* or *Desulfurococcales* to maintain or grow according to the presence or absence of an electron acceptor. These microorganisms are also recognized as the first colonizers found in newly formed chimneys [4, 5]. Thus, this source of electronic energy would be one of the easily accessible energy means for these electroautotrophic microorganisms on the same basis of molecular hydrogen, hydrogen sulfide or methane for chemotrophic microorganisms. To the same extent, electroautotrophic microorganisms enriched at the beginning of our experiment and using the reductive way of acetyl-CoA show that pyruvate would certainly be one of the central molecules in the development of the heterotrophic microbial biocenosis of these extreme ecosystems. Finally, this electrosynthesis of pyruvate by deep-branching *Archaea* through the ancient WL pathway could be compared to prebiotic organic synthesis from inorganic compounds involved in the origin of life theories in hydrothermal context. Pyruvate could so have been key metabolite for proto-metabolisms and be a building block for complex organic molecules synthesis. The constant production of electricity in the deep hydrothermal systems over time could have served as energetic driving force allowing the slow prebiotic synthesis reaction in a delimited space (organic micelles or mineral microcavity) to form the protometabolism and the protocell.

## Acknowledgments

This work received financial support from the CNRS national interdisciplinary research program (PEPS-ExoMod 2016). The authors thank Erwan Roussel (LM2E, IFREMER Brest) for helpful suggestions; Céline Rommevaux and Françoise Lesongeur for sampling during the MOMARSAT 2014 cruise; the MIM platform (MIO, France) for allowing providing access to their confocal microscopy facility; and the GeT-PlaGe platform (GenoToul, France) for DNA sequencing. The project leading to this publication has received funding from European FEDER Fund under project 1166-39417. The authors declare no conflict of interest.

## References

1. Kristall B, Kelley DS, Hannington MD, Delaney JR. Growth history of a diffusely venting sulfide structure from the Juan de Fuca Ridge: A petrological and geochemical study. Geochem Geophys Geosyst 2006; 7.

2. Kelley DS, Baross JA, Delaney JR. Volcanoes, fluids, and life at mid-ocean ridge spreading centers. Annu Rev Earth Planet Sci 2002; 30: 385–491.

3. Wirth R. Colonization of black smokers by hyperthermophilic microorganisms. Trends Microbiol 2017; 25: 92–99.

4. Pagé A, Margaret TK, Debra SS, Reysenbach A. Temporal and spatial archaeal colonization of hydrothermal vent deposits. Environ Microbiol 2008; 10: 874–884.

5. McCliment EA, Voglesonger KM, O’Day PA, Dunn EE, Holloway JR, Cary SC. Colonization of nascent, deep‐sea hydrothermal vents by a novel Archaeal and Nanoarchaeal assemblage. Environ Microbiol 2005; 8: 114–125.

6. Reysenbach A-L, Longnecker K, Kirshtein J. Novel bacterial and archaeal lineages from an in situ growth chamber deployed at a Mid-Atlantic Ridge hydrothermal vent. Appl Environ Microbiol 2000; 66: 3798–3806.

7. Flores GE, Campbell JH, Kirshtein JD, Meneghin J, Podar M, Steinberg JI, et al. Microbial community structure of hydrothermal deposits from geochemically different vent fields along the Mid-Atlantic Ridge. Environ Microbiol 2011; 13: 2158–2171.

8. Wirth R, Luckner M, Wanner G. Validation of a hypothesis: Colonization of black smokers by hyperthermophilic microorganisms. Front Microbiol 2018; 9: 524.

9. Yamamoto M, Nakamura R, Kasaya T, Kumagai H, Suzuki K, Takai K. Spontaneous and widespread electricity generation in natural deep-sea hydrothermal fields. Angew Chem Int Ed 2017; 56: 5725–5728.

10. Lovley DR. Electromicrobiology. Annu Rev Microbiol 2012; 66: 391–409.

11. Fu Q, Kuramochi Y, Fukushima N, Maeda H, Sato K, Kobayashi H. Bioelectrochemical analyses of the development of a thermophilic biocathode catalyzing electromethanogenesis. Environ Sci Technol 2015; 49: 1225–1232.

12. Fu Q, Kobayashi H, Kuramochi Y, Xu J, Wakayama T, Maeda H, et al. Bioelectrochemical analyses of a thermophilic biocathode catalyzing sustainable hydrogen production. Int J Hydrog Energy 2013; 38: 15638–15645.

13. Luo H, Teng W, Liu G, Zhang R, Lu Y. Sulfate reduction and microbial community of autotrophic biocathode in response to acidity. Process Biochem 2017; 54: 120–127.

14. He Z, Angenent LT. Application of bacterial biocathodes in microbial fuel cells. Electroanalysis 2006; 18: 2009–2015.

15. Aryal N, Tremblay P-L, Lizak DM, Zhang T. Performance of different Sporomusa species for the microbial electrosynthesis of acetate from carbon dioxide. Bioresour Technol 2017; 233: 184–190.

16. Beese-Vasbender PF, Grote J-P, Garrelfs J, Stratmann M, Mayrhofer KJJ. Selective microbial electrosynthesis of methane by a pure culture of a marine lithoautotrophic archaeon. Bioelectrochemistry 2015; 102: 50–55.

17. Pillot G, Davidson S, Shintu L, Ali OA, Godfroy A, Combet-Blanc Y, et al. Electrotrophy as potential primary metabolism for colonization of conductive surfaces in deep-sea hydrothermal chimneys. bioRxiv 2020; 2020.11.11.377697.

18. Wong ML, Charnay BD, Gao P, Yung YL, Russell MJ. Nitrogen oxides in early Earth’s atmosphere as electron acceptors for Life’s emergence. Astrobiology 2017; 17: 975–983.

19. Sarradin P-M, Cannat M. MOMARSAT2014 cruise, Pourquoi pas ? R/V. 2014. Sismer.

20. Pillot G, Davidson S, Auria R, Combet-Blanc Y, Godfroy A, Liebgott P-P. Production of Current by Syntrophy Between Exoelectrogenic and Fermentative Hyperthermophilic Microorganisms in Heterotrophic Biofilm from a Deep-Sea Hydrothermal Chimney. Microb Ecol 2019.

21. Pillot G, Frouin E, Pasero E, Godfroy A, Combet-Blanc Y, Davidson S, et al. Specific enrichment of hyperthermophilic electroactive *Archaea* from deep-sea hydrothermal vent on electrically conductive support. Bioresour Technol 2018; 259: 304–311.

22. Aminot A, Kérouel R. Dosage automatique des nutriments dans les eaux marines, 2007th ed. 2007. Ifremer.

23. Bax A, Davis DG. MLEV-17-based two-dimensional homonuclear magnetization transfer spectroscopy. J Magn Reson 1969 1985; 65: 355–360.

24. Schleucher J, Schwendinger M, Sattler M, Schmidt P, Schedletzky O, Glaser SJ, et al. A general enhancement scheme in heteronuclear multidimensional NMR employing pulsed field gradients. J Biomol NMR 1994; 4: 301–306.

25. Zhang L, Kang M, Xu J, Xu J, Shuai Y, Zhou X, et al. Bacterial and archaeal communities in the deep-sea sediments of inactive hydrothermal vents in the Southwest India Ridge. Sci Rep 2016; 6.

26. Bates ST, Berg-Lyons D, Caporaso JG, Walters WA, Knight R, Fierer N. Examining the global distribution of dominant archaeal populations in soil. ISME J 2011; 5: 908–917.

27. Rusch A, Amend JP. Order-specific 16S rRNA-targeted oligonucleotide probes for (hyper)thermophilic *Archaea* and *Bacteria*. Extremophiles 2004; 8: 357–366.

28. Teira E, Reinthaler T, Pernthaler A, Pernthaler J, Herndl GJ. Combining Catalyzed Reporter Deposition-Fluorescence In Situ Hybridization and Microautoradiography To Detect Substrate Utilization by Bacteria and Archaea in the Deep Ocean. Appl Environ Microbiol 2004; 70: 4411–4414.

29. Tamura K, Stecher G, Peterson D, Filipski A, Kumar S. MEGA6: Molecular Evolutionary Genetics Analysis Version 6.0. Mol Biol Evol 2013; 30: 2725–2729.

30. Yilmazel YD, Zhu X, Kim K-Y, Holmes DE, Logan BE. Electrical current generation in microbial electrolysis cells by hyperthermophilic archaea *Ferroglobus placidus* and *Geoglobus ahangari*. Bioelectrochemistry 2018; 119: 142–149.

31. Yee MO, Rotaru A-E. Extracellular electron uptake in Methanosarcinales is independent of multiheme c-type cytochromes. Sci Rep 2020; 10: 372.

32. Deutzmann JS, Sahin M, Spormann AM. Extracellular Enzymes Facilitate Electron Uptake in Biocorrosion and Bioelectrosynthesis. mBio 2015; 6: e00496–15.

33. Ishii T, Kawaichi S, Nakagawa H, Hashimoto K, Nakamura R. From chemolithoautotrophs to electrolithoautotrophs: CO2 fixation by Fe(II)-oxidizing bacteria coupled with direct uptake of electrons from solid electron sources. Front Microbiol 2015; 6.

34. Vorholt JA, Hafenbradl D, Stetter KO, Thauer RK. Pathways of autotrophic CO_2_ fixation and of dissimilatory nitrate reduction to N_2_O in *Ferroglobus placidus*. Arch Microbiol 1997; 167: 19–23.

35. Cabello P, Roldán MD, Moreno-Vivián C. Nitrate reduction and the nitrogen cycle in *Archaea*. Microbiology 2004; 150: 3527–3546.

36. Rosenberg E, DeLong EF, Lory S, Stackebrandt E, Thompson F (eds). The Prokaryotes: Other Major Lineages of Bacteria and The Archaea, 4th ed. 2014. Springer-Verlag, Berlin Heidelberg.

37. von Jan M, Lapidus A, Glavina Del Rio T, Copeland A, Tice H, Cheng J-F, et al. Complete genome sequence of Archaeoglobus profundus type strain (AV18T). Stand Genomic Sci 2010; 2: 327–346.

38. Schut GJ, Lipscomb GL, Han Y, Notey JS, Kelly RM, Adams MMW. The Order Thermococcales and the Family Thermococcaceae. In: Rosenberg E, DeLong EF, Lory S, Stackebrandt E, Thompson F (eds). The Prokaryotes. 2014. Springer Berlin Heidelberg, pp 363–383.

39. Chou C-J, Shockley KR, Conners SB, Lewis DL, Comfort DA, Adams MWW, et al. Impact of substrate glycoside linkage and elemental sulfur on bioenergetics of and hydrogen production by the hyperthermophilic archaeon *Pyrococcus furiosus*. Appl Environ Microbiol 2007; 73: 6842–6853.

40. Sokolova TG, Jeanthon C, Kostrikina NA, Chernyh NA, Lebedinsky AV, Stackebrandt E, et al. The first evidence of anaerobic CO oxidation coupled with H_2_ production by a hyperthermophilic archaeon isolated from a deep-sea hydrothermal vent. Extremophiles 2004; 8: 317–323.

41. Lee HS, Kang SG, Bae SS, Lim JK, Cho Y, Kim YJ, et al. The complete genome sequence of *Thermococcus onnurineus* NA1 reveals a mixed heterotrophic and carboxydotrophic metabolism. J Bacteriol 2008; 190: 7491–7499.

42. Seravalli J, Ragsdale SW. Channeling of carbon monoxide during anaerobic carbon dioxide fixation. Biochemistry 2000; 39: 1274–1277.

43. Oger P, Sokolova TG, Kozhevnikova DA, Chernyh NA, Bartlett DH, Bonch-Osmolovskaya EA, et al. Complete Genome Sequence of the Hyperthermophilic Archaeon Thermococcus sp. Strain AM4, Capable of Organotrophic Growth and Growth at the Expense of Hydrogenogenic or Sulfidogenic Oxidation of Carbon Monoxide. J Bacteriol 2011; 193: 7019–7020.

44. Bae SS, Kim TW, Lee HS, Kwon KK, Kim YJ, Kim M-S, et al. H2 production from CO, formate or starch using the hyperthermophilic archaeon, Thermococcusonnurineus. Biotechnol Lett 2012; 34: 75–79.

45. Huber H, Stetter KO. Desulfurococcales ord. nov. Bergey’s Manual of Systematics of Archaea and Bacteria. 2015. American Cancer Society, pp 1–2.

46. Albuquerque L, Costa MS da. The Family *Thermaceae*. The Prokaryotes. 2014. Springer, Berlin, Heidelberg, pp 955–987.

47. Borrel G, Adam PS, Gribaldo S. Methanogenesis and the Wood–Ljungdahl Pathway: An Ancient, Versatile, and Fragile Association. Genome Biol Evol 2016; 8: 1706–1711.

48. Berg IA, Kockelkorn D, Ramos-Vera WH, Say RF, Zarzycki J, Hügler M, et al. Autotrophic carbon fixation in archaea. Nat Rev Microbiol 2010; 8: 447–460.

49. Hocking WP, Stokke R, Roalkvam I, Steen IH. Identification of key components in the energy metabolism of the hyperthermophilic sulfate-reducing archaeon Archaeoglobus fulgidus by transcriptome analyses. Front Microbiol 2014; 5.

50. Utter M, Keech D. Pyruvate carboxylase. I. Nature of the reaction. J Biol Chem 1963; 238: 2603–2608.

51. Blamey JM, Adams MWW. Purification and characterization of pyruvate ferredoxin oxidoreductase from the hyperthermophilic archaeon *Pyrococcus furiosus*. Biochim Biophys Acta BBA - Protein Struct Mol Enzymol 1993; 1161: 19–27.

52. Huber H, Stetter KO. Desulfurococcales. The prokaryotes. 2006. Springer, pp 52–68.

53. Moon Y-J, Kwon J, Yun S-H, Lim HL, Kim M-S, Kang SG, et al. Proteome Analyses of Hydrogen-producing Hyperthermophilic Archaeon Thermococcus onnurineus NA1 in Different One-carbon Substrate Culture Conditions. Mol Cell Proteomics 2012; 11: M111.015420–M111.015420.

54. Lovley DR. Syntrophy goes electric: Direct interspecies electron transfer. Annu Rev Microbiol 2017; 71: 643–664.

55. Rotaru A-E, Shrestha PM, Liu F, Markovaite B, Chen S, Nevin KP, et al. Direct interspecies electron transfer between *Geobacter metallireducens* and *Methanosarcina barkeri*. Appl Environ Microbiol 2014; 80: 4599–4605.

56. Ward DE, Kengen SWM, van der Oost J, de Vos WM. Purification and Characterization of the Alanine Aminotransferase from the Hyperthermophilic Archaeon Pyrococcus furiosus and Its Role in Alanine Production. J Bacteriol 2000; 182: 2559–2566.

57. Brileya K, Reysenbach A-L. The Class *Archaeoglobi*. The Prokaryotes. 2014. Springer, Berlin, Heidelberg, pp 15–23.

